# Feasibility and repeatability of MEG-compatible patellar-tendon stimulator for eliciting knee-joint proprioceptive cortical responses

**DOI:** 10.64898/2026.06.26.734808

**Authors:** Feiyue Li, Anni Byman, Junru Chen, Toni Mujunen, Harri Piitulainen

**Affiliations:** Faculty of Sport and Health Sciences, University of Jyväskylä, Jyväskylä, Finland; Faculty of Educational Sciences, University of Helsinki, Helsinki, Finland; Center for Interdisciplinary Brain Research, University of Jyväskylä, Jyväskylä, Finland; Center for Neuroplasticity and Pain, Department of Health Science and Technology, Faculty of Medicine, Aalborg University, Aalborg, Denmark

**Keywords:** MEG-compatible stimulator, Patellar tendon reflex, Repeatability, Movement evoked field, Beta rhythm, Somatosensory, Proprioception

## Abstract

**Background:** Cortical processing of the knee-joint proprioception is largely unknown. Magnetoencephalography (MEG) can be used to quantify the cortical processing of the proprioceptive afference, but MEG-compatible and well-controlled stimulation of the knee joint is technically challenging, and thus has received less attention.

**New method:** We introduced a novel MEG-compatible stimulator that delivers controlled patellar tendon stretches to activate muscle afferent of the knee extensors. The stimulus intensity is adjustable, allowing graded activation of proprioceptive input and, when required, elicitation of the patellar-tendon reflex.

**Results:** The novel stimulator elicited clear muscular and cortical responses in both intensity conditions. Cortical responses demonstrated moderate to excellent intersession reliability for peak evoked field amplitude (ICC: 0.69–0.96), beta suppression (0.89–0.90) and beta rebound (0.96–0.97). Notably, beta suppression peaked more laterally than expected in both hemispheres. Peak EMG amplitudes in VL and VM muscles were reliable for both intensity conditions (ICC: 0.66–0.89), and stimulus kinematics remained consistent throughout measurements.

**Comparison with existing methods:** Previous robotic or motor-driven devices have been used to evoke cortical responses to knee-joint proprioceptive stimulation, but mechanical coupling across adjacent joints may limit knee-specific input. The present stimulator provides mechanically simple and MEG-compatible alternative that targets knee extensor afferents more directly, reduces distal joint involvement.

**Conclusion:** The novel stimulator is a feasible and repeatable tool to study cortical processing of proprioceptive afference from the knee-joint using MEG. The spatially unexpected beta rhythm suppression suggests that knee-joint proprioceptive afference may involve more unique sensorimotor cortical neuronal network than previously recognized.

## 1. Introduction

Evoked and induced cortical responses, as well as coherence between kinematics and cortical activity (Bourguignon et al., 2011; Bourguignon et al., 2015; Piitulainen et al., 2018b), can be measured with magnetoencephalography (MEG) or electroencephalography (EEG), and thus have been used to quantify cortical processing of proprioceptive afference during passive movements (Alary et al., 2002; Druschky et al., 2003; Lange et al., 2001). Evoked fields are time and phase-locked to the stimulus, whereas induced responses are not, but reflect activation and inhibition of the primary sensorimotor (SM1) cortex (David et al., 2006; Lakatos et al., 2009). Prominent ∼20-Hz beta rhythm or activity is observed in the SM1 cortex, and its power has been shown to modulate following somatosensory stimuli or motor activities (Gaetz and Cheyne, 2006; Houdayer et al., 2006; Laaksonen et al., 2012; Parkkonen et al., 2015; Salmelin and Hari, 1994). Typically, beta power in the SM1 cortex is suppressed after somatosensory stimulation (suppression, or event-related desynchronization), which is thought to reflect the activation of the SM1 cortex (Neuper et al., 2006; Pfurtscheller, 2001). The suppression is followed by a delayed rebound of beta power above its baseline level (rebound, or event-related synchronization). The rebound is associated with cortical deactivation or inhibition of the SM1 cortex (Neuper et al., 2006; Pfurtscheller, 2001). Anatomically, the beta suppression and rebound have different spatial distribution in the SM1 cortex, for example, the beta rebound is located more anterior than the suppression (Cheyne et al., 2006; Jurkiewicz et al., 2006). The beta rhythm is also altered by aging (Rossiter et al., 2014b) and diseases, such as Parkinson’s disease (Heinrichs-Graham et al., 2014; Vinding et al., 2019) and stroke (Rossiter et al., 2014a).

Cortical processing of proprioceptive afference arising from the hand has been quantified using MEG and EEG, particularly during active or passive finger movement (Illman et al., 2022; Illman et al., 2020; Lange et al., 2001; Piitulainen et al., 2013). In contrast, the lower-limb proprioception has received less attention, with most previous work focusing on the ankle joint (Mujunen et al., 2022; Piitulainen et al., 2018b) rather than the knee. The knee joint plays a central role in maintaining postural control, motor coordination, and joint stability (Solomonow and Krogsgaard, 2001). Proprioceptive afference from the knee joint arises from proprioceptors, located in muscles, ligaments, and joint capsules, which together contribute to the sensorimotor integration required for stable and efficient movement (Banios et al., 2022; Daneshjoo et al., 2012). However, previous studies have reported that proprioceptive abilities of the knee to sense joint position and joint motion are impaired with traumatic injuries, pathological conditions, and age-related degeneration (Barrett et al., 1991; Koralewicz and Engh, 2000; Nyland et al., 2017). Accordingly, it is unclear whether, and to what extent, the cortical processing of proprioceptive afference from the knee is altered in various conditions.

Different robotic or motor-driven devices have been used to elicit cortical responses to proprioceptive stimulation of the knee joint in functional magnetic resonance imaging (fMRI) or EEG during the past two decades (Hollnagel et al., 2011; Jaeger et al., 2014; Martinez et al., 2014; Zhao et al., 2025). However, isolated stimulation of the knee joint has proved to be challenging due to the knee joint anatomical structure. For example, the stepping robot (Hollnagel et al., 2011; Martinez et al., 2014) or similar mechanical approaches (Zhao et al., 2025) may unintentionally activate the proprioceptors of the ankle joint as well, thereby limiting the specific stimulation of the knee joint. In addition, it is noteworthy that MEG signals are highly sensitive to head movement, especially in motor task studies. Therefore, the knee stimulator must be carefully designed to minimize unintended body and head movements while ensuring full MEG compatibility. This can be achieved through stable leg fixation, controlled stimulus delivery, and the use of non-magnetic components.

Here we introduce a novel MEG-compatible stimulator to briefly stretch to the patellar tendon and thus activating the muscle afferents of the knee extensors. By adjusting the stimulus intensity, the patellar-tendon reflex can also be evoked, and the resulting muscle activity can be quantified using electromyography (EMG). The patellar-tendon reflex is a classical monosynaptic stretch reflex involving contraction of the quadriceps muscles in response to the sudden patellar tendon stretch.

We aimed to test feasibility and repeatability of the novel MEG-compatible stimulator in eliciting cortical responses to knee-joint proprioceptive afference. We tested both high and low intensity conditions to clarify whether repeatability is dependent on the stimulation intensity or not. Based on previous studies demonstrating a good to excellent reproducibility of the novel finger or ankle stimulator in MEG (Chen et al., 2025; Illman et al., 2022; Mujunen et al., 2022; Piitulainen et al., 2015; Piitulainen et al., 2018b), we hypothesized that the stimulator would generate reliable proprioceptive responses across sessions, making it suitable for studying cortical proprioceptive processing of the knee-joint with MEG.

## 2. Materials and methods

### 2.1 Subjects

In total twenty-five healthy participants (25 right-foot dominant, the footedness score: 9.3 ± 4.2; mean ± SD, age: 30.3 ± 4.3yr; 11 females) were recruited in this study, but only twenty-three participants (age: 29.5 ± 2 yr, 11 females) completed both measurement sessions. The footedness was assessed by the Waterloo footedness questionnaire (Elias et al., 1998). The participants gave their written informed consent before the measurements. The procedures of the study were approved by the University of Jyväskylä ethics committee (694/13.00.04.00/2023) and conducted in accordance with the Declaration of Helsinki.

### 2.2 MEG-compatible knee joint stimulator

Fig. 1 shows the custom-made MEG-compatible stimulator to elicit proprioceptive stimulation through patellar tendon taps using POM-plastic hammer during MEG recordings. The hammer was supported by a PVC-plastic frame surrounding a modified MEG chair that enables force recordings. The chair was a commercial MEG chair (Elekta Oy, Helsinki, Finland) modified by the Faculty of Sport and Health Science (University of Jyväskylä, Finland) to integrate non-magnetic strain gauges into aluminum plates, which were affixed to the chair’s leg supports. The stimulator frame included: the main framework, horizontal guide rods and stimulus-height controller. The guide rods were vertically and horizontally adjustable to accommodate different leg lengths and allow precise lateral positioning of the hammer for accurate targeting of the stimulation site. Stimulus-height controller allows fine-control of hammer’s drop height to adjust the stimulation intensity. The stimulus height can be further elevated by placing plastic panels underneath the device, e.g., in case of short individuals or child recordings.

**Fig. 1.**
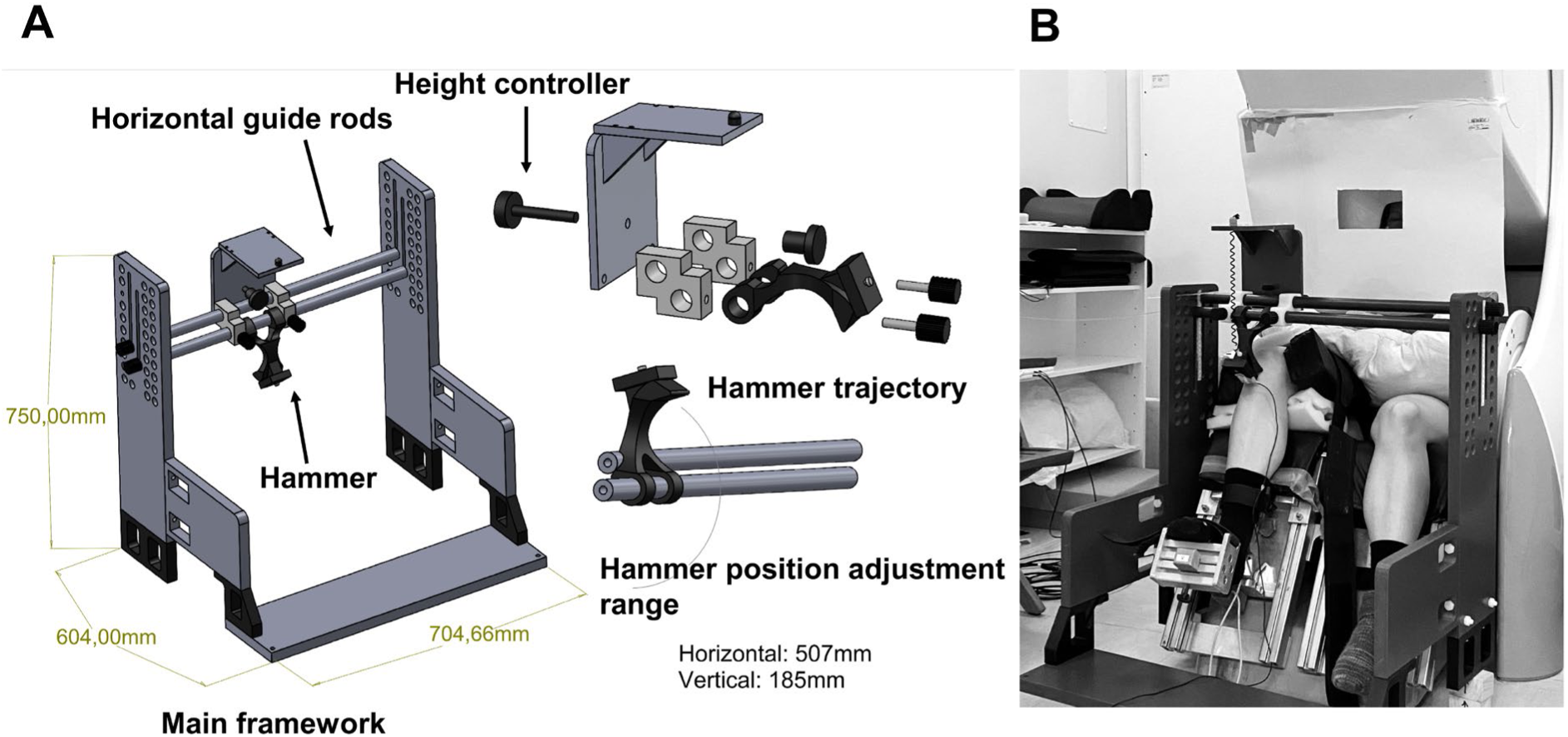
Technical drawing of the MEG-compatible knee joint stimulator used to elicit patellar tendon reflex. **A.** All materials and components are plastic and thus nonmagnetic. **B.** The knee-joint stimulator in the MEG lab.

The hammer was first placed to the initial “up” position by the experimenter. After visual cue, the experimenter released the hammer, allowing it to fall in a gravity-driven arc toward the patellar tendon. The experimenter then captured the hammer immediately when it bounced back, and returned it back to the initial “up” position to prepare for the next stimulus.

### 2.3 Experimental protocol

In this study, two separate measurements (separated by 46 ± 15 days, range: 31−61 days) were conducted at the Centre for Interdisciplinary Brain Research, University of Jyväskylä. During the recording, the participant was instructed to sit and remain relaxed and not pay attention to the stimuli (Fig. 1B). The participants’ arms were positioned on top of a pillow placed on their lap, while the participant’s right foot was secured against the pedal-like foot support of the custom-made MEG chair with a Velcro belt. Foam padding and adjustable straps were used around the right thigh and shin to ensure comfort and to minimize motion artefacts. A cardboard, taped in front of the participant, blocked visual contamination from the reflex hammer, the knee joint and the experimenter operating the hammer. The participant was asked to fixate on a marker placed in front of the wall of the magnetically shielded room. To limit head movements during the MEG recordings, pieces of foam cushion were placed between the participant’s forehead and MEG-helmet and secured with a net cap. In addition, to block any minor auditory disturbances arising from the reflex hammer, participants wore earplugs, and Brownian noise was played through flat panel speakers. One experimenter remained inside the shielded room with the participant during the proprioceptive stimulation to operate the stimulation device.

Fig. 2A shows the proprioceptive stimuli at two different intensities, high and low, to the patellar tendon with inter-stimulus interval (ISI) of 5.0−6.0 s (mean ∼5.5 s). The stimulus timing was indicated to the experimenter through light signal by optical fiber using Presentation software (Ver. 21.1, Neurobehavioral Systems Inc., Albany, CA, USA). A total of 100 stimuli were delivered for each intensity condition. The stimulation intensity for the first session was randomized for the first participant and then alternated for the following participants, i.e., if the previous participant started with the high intensity condition, then the next participant started with the low intensity condition. The order of the intensity condition within each individual participant was kept the same between the first and second measurements, while the recording time was also at the same time of the day for each individual.

**Fig. 2.**
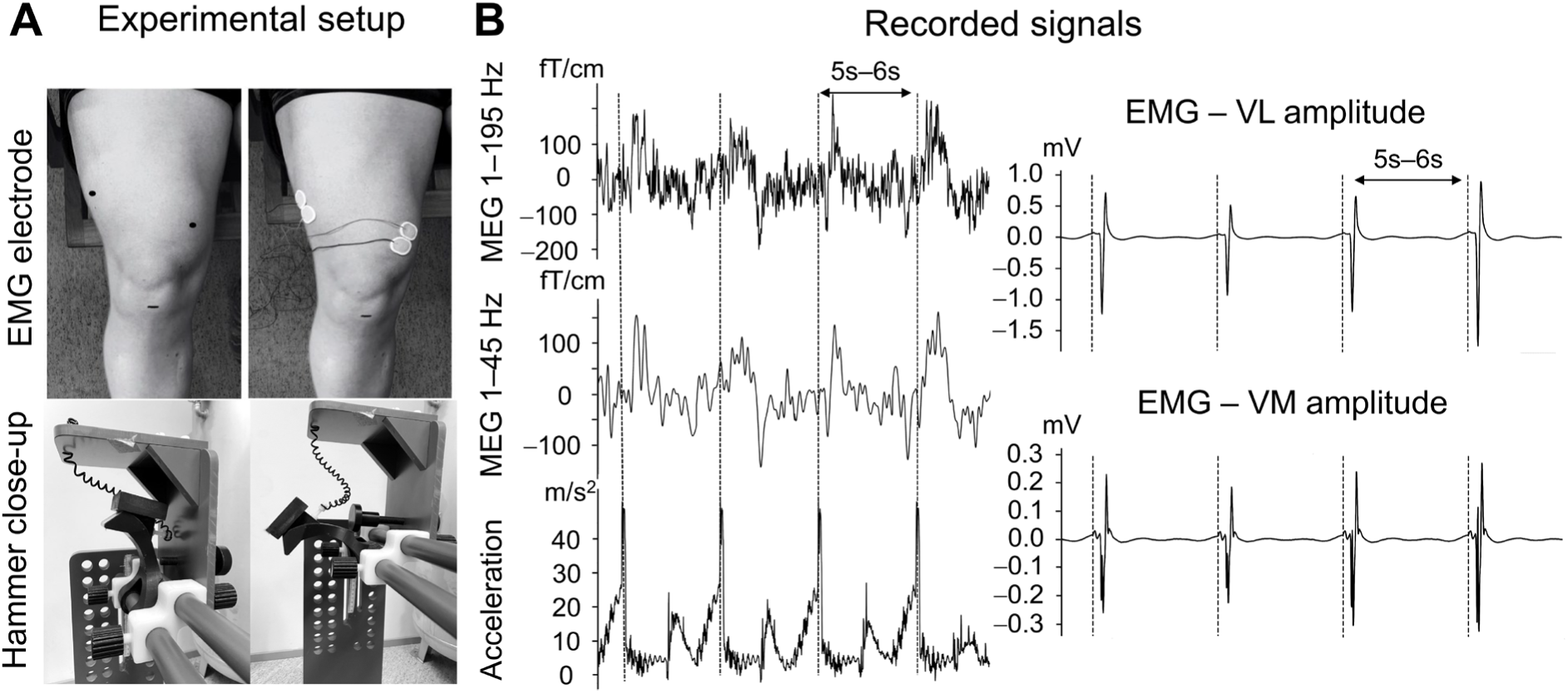
Experimental setup and representative signals. **A.** Placement of the EMG electrodes on top of the vastus lateralis (VL) and vastus medialis (VM) muscles, and the reference point of the patellar tendon stimulation location (top panels). The hammer positions for high intensity condition (bottom left panel) and low intensity condition (bottom right panel). **B.** MEG signals with 1−195 Hz and 1−45 Hz from one representative channel, acceleration magnitude and EMG amplitude in a representative participant. The grey vertical dashed lines indicate the stimulus onsets.

### 2.4 Signal acquisition

#### Cortical MEG activity and MRI

MEG signals were acquired in a magnetically shielded room using a 306-channel whole-scalp neuromagnetometer (Elekta Neuromag TRIUXTM, Elekta Oy, Helsinki, Finland). MEG signal sampling rate was 1000 Hz with a 0.1−330 Hz bandpass filter. The eye blink artefacts were recorded with electro-oculography, including two electrodes at 1 cm lateral to the left and right outer canthi. The ground electrode was placed on top of the right clavicle. Before data acquisition, five head position coils were attached to the scalp to continuously monitor the participant’s head position with respect to the MEG sensors during the recording. The location of three anatomical landmarks, five head position coils and additional points (over ∼200 points) on the head surface were digitized by using a 3-D digitizer (Isotrak, Polhemus, Colchester, VT, USA). The head position and the quality of all recording signals were checked at the beginning of each recording and recorded continuously during the measurement.

Structural MRI data was not collected in the present experiment. However, anatomical MRIs were acquired for four of our participants in another recent study. GE 1.5T Signa HDxt MRI scanner (GE Healthcare, Milwaukee, WI, USA) with a standard head coil was used to scan T1- and T2-weighted MRIs of the head in Synlab Oy (Jyväskylä, Finland). Then these MRIs were used in source-level analysis of the MEG data.

#### Muscular responses and stimulus kinematics

The surface EMG and accelerations were recorded time-locked to MEG signals with the same sampling frequency of 1000 Hz with 0.1−330 Hz passband. To determine stimulation onsets and quantify the stimulus kinematics, acceleration of the reflex hammer was recorded with a 3-axis accelerometer (ADXL335 iMEMS Accelerometer, Analog Devices Inc. Norwood, MA, USA) attached to the top of the hammer.

The stretch reflex responses of quadriceps muscles were recorded with two pairs of Ag/AgCl electrodes (Ambu Neuroline 720 15-K/C/12, Denmark) taped on the vastus lateralis (VL) and vastus medialis (VM) muscles (Fig. 2A). Prior to placing the EMG electrode, body hair and dead skin were removed with razor and a piece of sandpaper, respectively. We aimed to avoid innervation zone regions as recommended by Barbero et al. (2012): VL at between 0 and 43 mm on the line on the distal portion of the muscle belly and oriented 20° with respect to the reference line between the lateral side of the patella and the anterior superior iliac spine, and VM at between 0 and 38 mm on the line on the distal portion of the muscle belly and oriented 50° with respect to the reference line between the medial side of the patella and the anterior superior iliac spine. The quality of EMG and accelerations signals were confirmed based on the raw data before starting the actual experimental recordings.

### 2.5 Data processing and analysis

#### MEG preprocessing

All MEG raw signals were first visually checked to select noisy MEG sensors. Next, a custom-made script using MNE-Python (Gramfort et al., 2014) was used to obtain and average the head position coordinates across different days (Day-1, Day-2) and intensity conditions (high intensity, low intensity). The average head position was used to compensate for head movement as the MEG signals were preprocessed with MaxFilter software (v. 3.0, Elekta Neuromag Oy, Helsinki, Finland) using the signal-space separation method with temporal extension (tSSS) to reduce the external interference (Taulu and Simola, 2006). Lastly, an independent component analysis implemented in MNE-Python was performed to remove eye blink artifacts and cardiac activity based on 30 decomposed components from MEG signals (Hyvarinen and Oja, 2000), which were band-pass filtered at 1–40 Hz using a zero-phase finite impulse response filter (firwin in SciPy; Hamming window).

#### Muscular responses and stimulus kinematics

Stimulation onsets were determined from the Euclidian norm (i.e., acceleration magnitude signal) of the three orthogonal acceleration signals bandpass filtered to 1–400 Hz. Stimulus onset (i.e., hammer percussion) was determined as the time corresponding to the peak acceleration magnitude separately for each participant, intensity condition, and day.

EMG signals were band-pass filtered at 10–295 Hz and epoched from –1000 to 1000 ms around the stimulus onset at 0 ms. After the epoching, the peak-to-peak amplitude of the muscular EMG responses were calculated. The stimulus-to-stimulus variations of the VL and VM muscles were also computed for each participant.

#### Cortical MEG responses − evoked field

Preprocessed MEG signals were first band-pass filtered at 1−95 Hz, after which the data were epoched with a time window of −2000 to 2000 ms with respect to the stimulus onset. During the epoching, the individual epoch was excluded if the signal amplitude exceeded 4 pT for magnetometers and 4 pT/cm for gradiometers. A total of 99 ± 3 (high intensity) and 98 ± 3 (low intensity) epochs for Day-1, and 97 ± 6 and 97 ± 5 epochs for Day-2, were retained and averaged for the final analysis. Two gradiometer signals for each pair were combined by calculating the vector sum to replace magnetometer signals for the whole helmet. The gradiometer pair from the parietal sensors displaying the largest evoked field amplitude was chosen as the peak gradiometer pair for the final analysis. The peak gradiometer pair was selected independently in each day and intensity condition, with some participants showing the same peak gradiometer pair across days and intensity conditions.

#### Cortical MEG responses − induced response

To quantify the strength of the stimulus-related beta rhythm modulation, the temporal spectral evolution (TSE) method was used in all intensity conditions and days (Salmelin and Hari, 1994). Preprocessed MEG signals were first band-pass filtered with a beta frequency band 12−31 Hz and epoched from −2000 to 3000 ms with respect to the stimulus onset. The evoked fields were subtracted from the data (David et al., 2006), after which the data were rectified and averaged across epochs in both conditions and days for further analysis. Peak strengths and latencies of the beta rhythm were individually determined from the TSE curves. The peak gradiometer, demonstrating the most robust beta suppression and rebound, was defined separately for each participant, intensity condition, and day. In some participants, the rebound and suppression of beta rhythm were more pronounced in different gradiometers. If there were no visible beta rhythm rebound or suppression in one day or one intensity condition, the participant was excluded from the analysis. The baseline beta rhythm power was computed from the pre-stimulus time window (from −1800 to −100 ms). The absolute suppression and rebound strengths were calculated and converted to the relative value (in percentage) with respect to the baseline.

#### Source-level analysis of the induced beta suppression and rebound

We performed induced response analysis at the source-level for a sub-group of the participants (n = 4; all who had MRIs available) to clarify the unexpected spatial distribution (peaking in lateral to the hand area) of the beta suppression. The analysis was performed identically across days and conditions. First, cortical surface reconstruction was performed using FreeSurfer’s recon-all pipeline on the original T1-and T2-weighted anatomical images. Following reconstruction, the cortical and skull surfaces were visually inspected for possible segmentation errors. If residual skull or dura were present, it was manually removed prior to boundary element model (BEM) creation. A single-compartment BEM of the inner skull was generated using FreeSurfer’s watershed algorithm.

During the MEG preprocessing, the same procedure described above in the sensor-level analysis was applied, with the exception that the head position coordinates were kept day and intensity condition specific. Then, MEG-MRI co-registration was performed using digitized fiducial landmarks (nasion and left/right preauricular points) and additional head shape points. An automated co-registration procedure implemented in MNE-Python was used, and the resulting alignment was visually inspected and manually adjusted when necessary. The forward solution was computed using the BEM model, co-registration matrix, and a surface-based source space.

Preprocessed MEG signals were band-pass filtered in the beta frequency range (12–31 Hz) and epoched from −2000 to 3000 ms relative to the stimulus onset. The evoked field was subtracted from the epoched data. An inverse operator was then created from the forward solution for source estimation. After inversing, the beta power was estimated using the source_induced_power function in MNE-Python, applying the relative activity (in percentage) of the baseline period (−1800 ms to −100 ms) and exact low-resolution brain electromagnetic tomography (eLORETA) across the whole source space (Pascual-Marqui et al., 2011). Finally, to characterize the spatial distribution of cortical induced responses to proprioceptive stimulation, the contralateral (left) hemisphere to the stimulated knee was selected as the region of interest for further analysis.

### 2.6 Statistical analysis

Statistical analysis was performed in IBM SPSS software (v. 27.0). The normality of the data was assessed using Shapiro-Wilk test. The correlations and intraclass correlation coefficients (ICC, model 3,1; two-way mixed-effects, absolute agreement) were calculated to assess the reliability of kinematic, muscular, evoked field, and induced response parameters between days. ICC values of < 0.4 were indicative of poor, 0.5−0.75 as moderate, 0.75−0.9 as good and >0.9 as excellent reliability (Koo and Li, 2016). In addition, coefficient of variation (CoV) was defined to assess inter-session variability at each condition. The inter-session variation was defined as the ratio of the standard deviation to the mean (CoV = SD / mean × 100%).

Due to the non-normal distribution of the latencies of induced responses (beta rebound and suppression), the nonparametric Wilcoxon test was used to test the inter-session differences. Lastly, one-way repeated measures multivariate analysis of variance (MANOVA) was conducted to examine differences for normally distributed variables, including the head position (x, y and z axis), peak acceleration magnitude, EMG responses, evoked-field amplitude, and the relative peak strengths of beta rebound and suppression between days.

## 3. Results

Fig. 2B shows the recorded MEG, EMG and acceleration signals from a representative participant during high intensity condition at Day-1. The MEG cortical responses are visible even from individual raw-signal responses.

From twenty-three participants, four were excluded from the final MEG analyses due to low number of epochs per intensity condition (below 80 out of 100 epochs) and other technical difficulties. Totally, nineteen participants (mean ± SD, age: 28.6 ± 2.2 yr; 10 females) were included in the further analysis. For the induced analysis, one participant was excluded, since the modulation of the ∼20-Hz power was below the background level. This is typical for some individuals when quantifying rhythmic modulations of the SM1 cortex for the proprioceptive stimulation of the lower limb (Giangrande et al., 2024; Mujunen et al., 2022; Mujunen et al., 2025). For the EMG analysis, one participant was excluded due to the lack of visible and clear stretch-reflex response.

A repeated-measures MANOVA with session as within-subjects factor across the above 20 dependent variables did not reveal statistically significant differences (_F20, 15_ = 0.86, p = 0.63). Note that although some variables had 19 valid cases and others 18, the MANOVA requires complete data across all included variables. Therefore, only participants with complete datasets across all 20 variables (n = 18) were retained in the multivariate test.

### 3.1 Head position and stimulus kinematics

Inter-session difference in the head position within MEG helmet was only few millimeters, and no statistically significant differences were observed between Day-1 and Day-2 in both intensity conditions (p > 0.05). The absolute differences in head position between Day-1 and Day-2 were 0.80 ± 2.35 mm for x-axis, 2.10 ± 5.41 mm for y-axis and 1.21 ± 3.01 mm for z-axis in high intensity condition, and 0.60 ± 2.59 mm for x-axis, 1.24 ± 4.50 mm for y-axis, and 1.60 ± 3.70 mm for z-axis in low intensity condition.

The top panels of Fig. 3A show group averaged acceleration curves across subjects (n = 19) for both intensity conditions and days. Mean values and standard deviations of the peak acceleration magnitude for both intensity conditions on two days are shown in Table 1. The group level inter-session differences were 4.7 ± 5.1 m/s^2^ (0−16.1 m/s^2^) for high intensity condition and 3.1 ± 4.0 m/s^2^ (0−16.4 m/s^2^) for low intensity condition, with no statistically significant difference between days (p > 0.05).

**Fig. 3.**
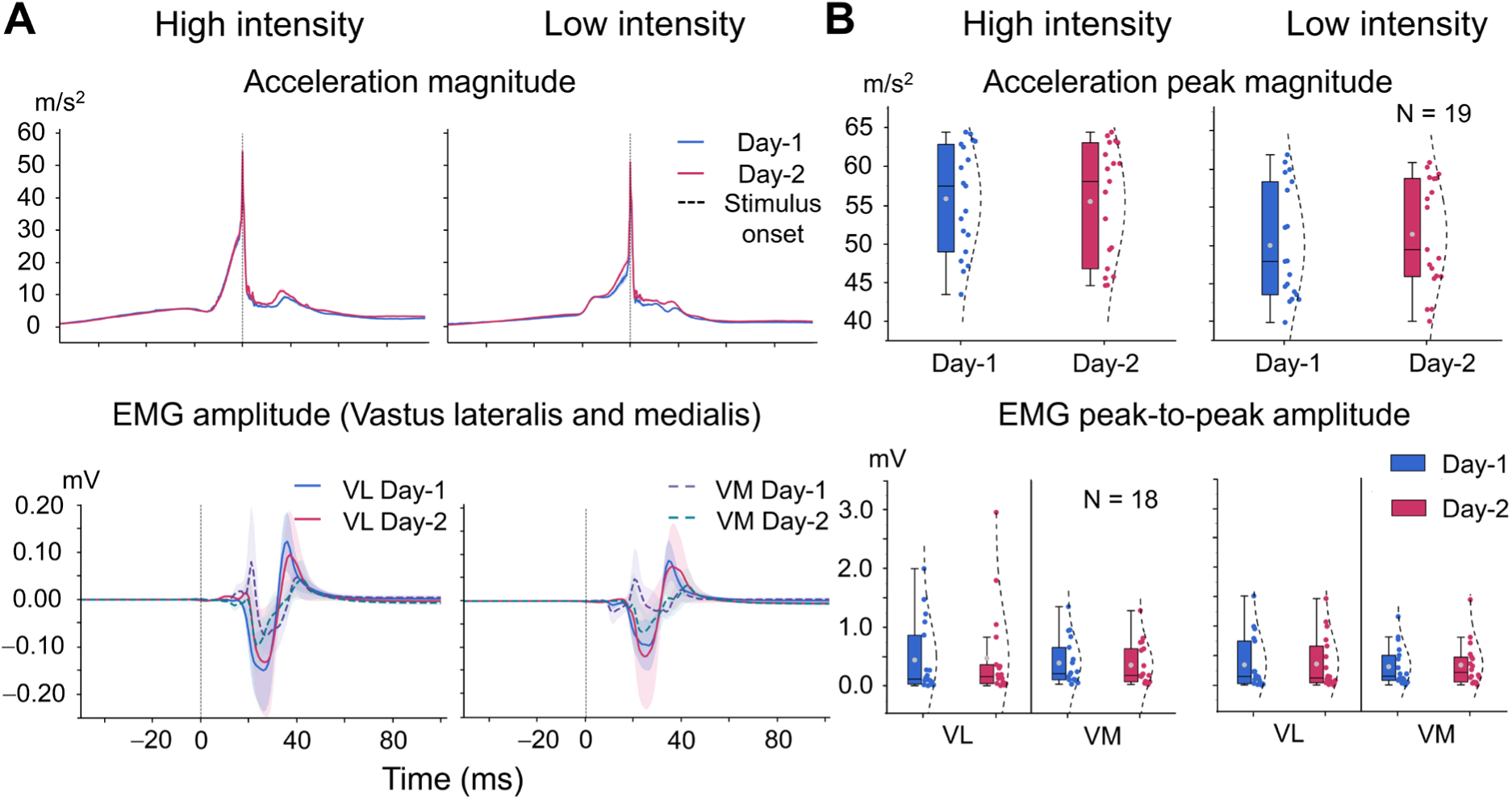
Acceleration magnitude and muscular responses under high and low intensity conditions at Day-1 and Day-2. **A.** Top panels show grand average acceleration magnitude curves with standard deviations (shade area). The bottom panels show pooled grand average EMG amplitude curves with standard deviations. The grey vertical dashed lines indicate the stimulus onset. **B.** The acceleration peak magnitude (top) and the peak-to-peak amplitudes of all muscles (bottom) for both days and intensity conditions. In the box plots, the boxes include fifty percent of peak values and horizontal lines in boxes indicate median values. The points (blue and red colors) in dot plots indicate the individual values and overlaid dashed curves represent the distribution of values. The whiskers show the range of data.

**Table 1.** Pooled kinematics, muscular, evoked and induced response parameters (mean ± SD) for both conditions in the two measurement days.

|  | High intensity |  | Low intensity |  |
| --- | --- | --- | --- | --- |
|  | Day-1 | Day-2 | Day-1 | Day-2 |
| **Kinematics** |  |  |  |  |
| Peak magnitude, $\text{m/s}^2$ | $55.8 \pm 7.0$ | $55.5 \pm 7.5$ | $50.0 \pm 7.4$ | $51.4 \pm 7.5$ |
| Range | 44.6–64.2 | 44.6–64.4 | 42.6–61.9 | 40.0–60.9 |
| **EMG** |  |  |  |  |
| VL amplitude, mV | $0.45 \pm 0.61$ | $0.54 \pm 0.85$ | $0.35 \pm 0.45$ | $0.47 \pm 0.65$ |
| Range | 0.02–2.01 | 0.02–2.96 | 0.01–1.51 | 0.01–2.00 |
| VM amplitude, mV | $0.40 \pm 0.39$ | $0.37 \pm 0.36$ | $0.32 \pm 0.33$ | $0.34 \pm 0.37$ |
| Range | 0.04–1.36 | 0.04–1.29 | 0.01–1.12 | 0.01–1.45 |
| **Evoked field** |  |  |  |  |
| Peak evoked amplitude, fT/cm | $160.8 \pm 49.9$ | $158.5 \pm 44.5$ | $132.7 \pm 53.5$ | $137.7 \pm 40.9$ |
| Range | 58.9–269.8 | 48.9–241.8 | 29.5–244.7 | 55.6–197.7 |
| Latency, ms | $54 \pm 24$ | $54 \pm 25$ | $57 \pm 24$ | $58 \pm 23$ |
| **Induced response** |  |  |  |  |
| Relative suppression amplitude (%) | $-31.3 \pm 7.9$ | $-31.6 \pm 8.0$ | $-26.4 \pm 8.8$ | $-24.6 \pm 9.0$ |
| Range | –(16.6–49.6) | –(17.0–47.7) | –(14.3–40.8) | –(16.6–39.9) |
| Latency, ms | $293 \pm 78$ | $271 \pm 81$ | $221 \pm 54$ | $229 \pm 76$ |
| Relative rebound amplitude (%) | $55.2 \pm 24.8$ | $56.1 \pm 23.1$ | $45.0 \pm 19.4$ | $45.4 \pm 19.9$ |
| Range | 20.4–94.2 | 27.5–95.5 | 20.5–77.4 | 16.7–83.7 |
| Latency, ms | $751 \pm 111$ | $752 \pm 112$ | $748 \pm 117$ | $762 \pm 119$ |

VL: vastus lateralis muscle; VM: vastus medialis muscle.

### 3.2 Muscular response strength

The bottom panels of Fig. 3 illustrate EMG responses and their peak-to-peak amplitude for VL and VM muscles at Day-1 and Day-2 under both high and low intensity conditions. The grand average EMG amplitude curves of VL and VM muscles appear similar between Day-1 and Day-2, while it is greater in VL muscle than in VM muscle on both days and intensity conditions. Mean peak-to-peak EMG amplitudes of VL and VM muscles for both intensity conditions are shown in Table 1. No statistically significant differences were observed between different days in either intensity condition (p > 0.05).

### 3.3 Cortical evoked field strength

The top panels of Fig. 4 illustrate the group averaged time-series of the evoked field and peak evoked field amplitude for both intensity conditions across days. Both conditions evoked responses which were well identifiable on both days, and the curves of the evoked fields appear similar between Day-1 and Day-2, as left panel shows. The evoked field was most pronounced at around 54 ms and 57 ms after the onset of high and low intensity stimuli, respectively. All results for peak evoked field amplitudes and latencies are shown in Table 1. Peak evoked field amplitudes did not differ statistically significantly between the two days in either intensity condition (p > 0.05).

**Fig. 4.**
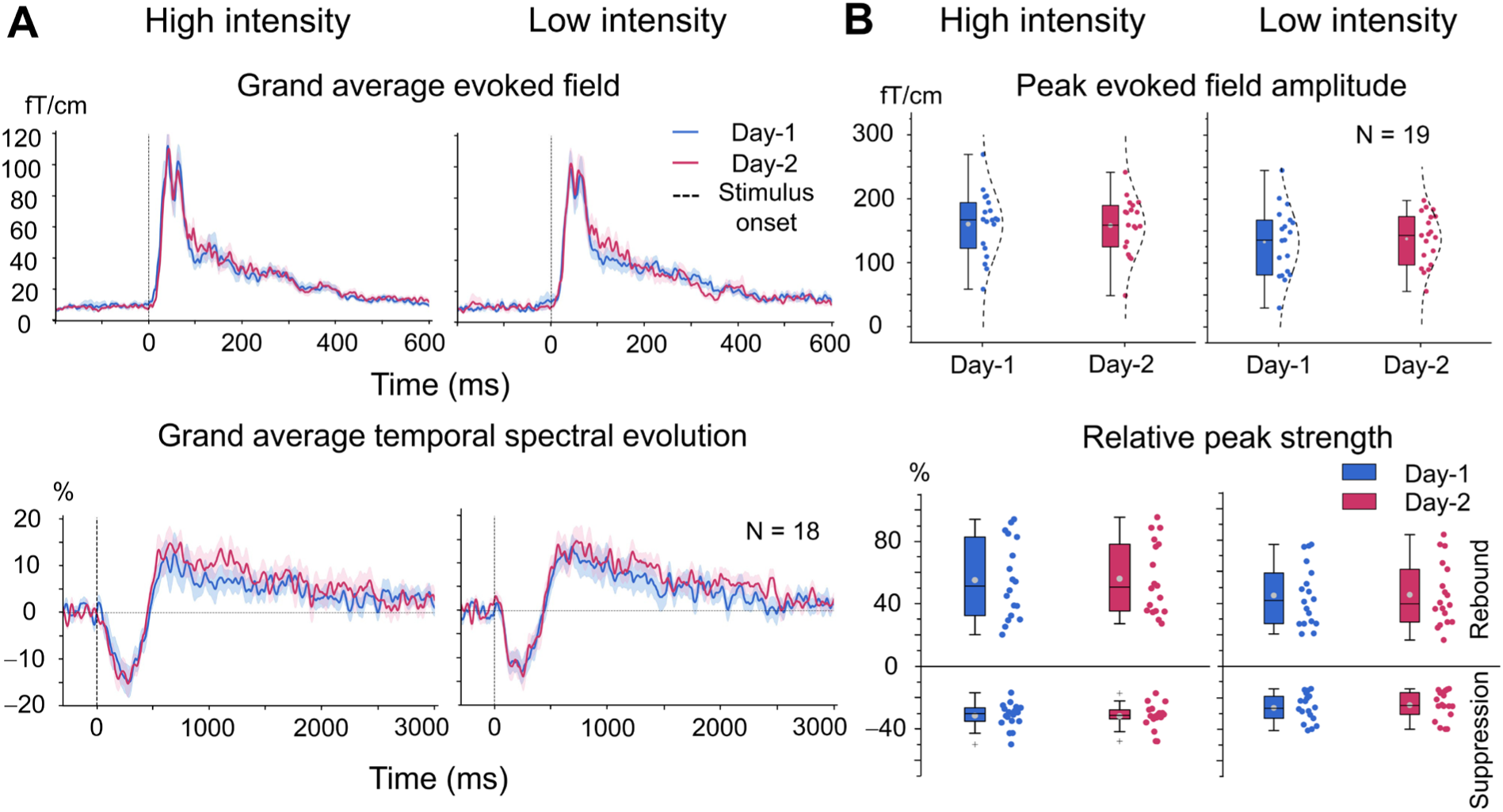
The cortical response strengths to the proprioceptive stimulation. **A.** Grand averaged (n = 19) time-series curves of evoked fields (the top panels) and temporal spectral evolution (TSE) time-series (n = 18) from one most representative channel over the left primary sensorimotor area (the bottom panels) for high intensity and low intensity conditions during Day-1 and Day-2. Grey vertical dashed lines indicate the onset of the stimulus at time zero. **B.** The values of peak evoked field amplitudes (top) and relative peak strengths of beta suppression and rebound (bottom) in high and low intensity conditions. Peak relative strength of beta rhythm suppression and rebound separately are shown as box plots across days in both intensity conditions.

### 3.4 Cortical induced response strength

The bottom panels of Fig. 4A show group level averaged TSE curves to high and low intensity conditions on both days. Both intensity conditions induced the well identifiable beta suppression and rebound on both days, with similar group averaged TSE curve shapes observed between Day-1 and Day-2 in both intensity conditions. The relative peak strengths and latencies of beta rhythm suppression and rebound are shown in Table 1. In addition, no statistically significant differences were observed in the relative peak strengths and latencies of beta suppression or rebound between Day-1 and Day-2 for either intensity conditions (p > 0.05).

### 3.5 Spatial distribution of the cortical responses

Fig. 5A shows the group averaged (n = 19) topographic distribution of the evoked field to high and low intensity conditions on both days. Peak evoked fields were observed over the contralateral leg region of SM1 cortex shortly after the onset of both high and low intensity conditions.

**Fig. 5.**
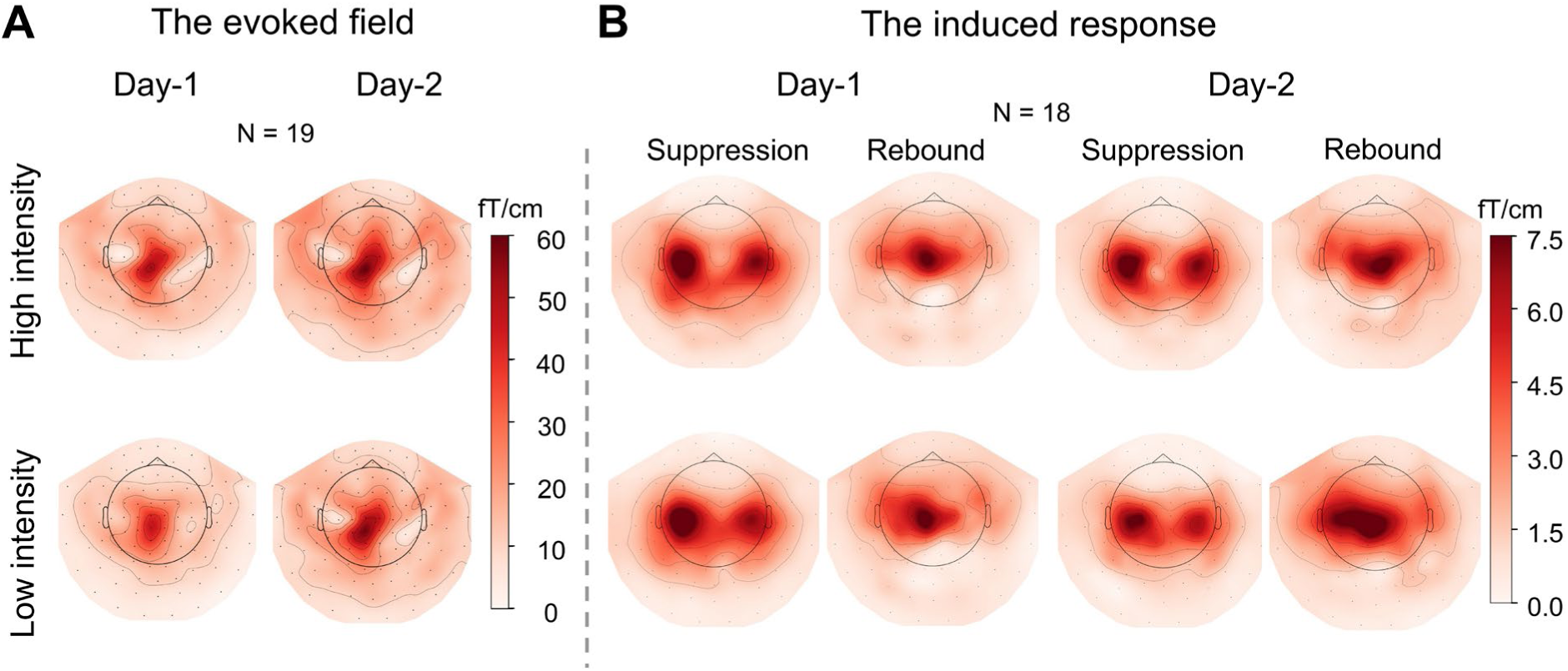
The cortical spatial distribution to proprioceptive stimulation. **A.** Grand average spatial distribution of the evoked field (n = 19) to high and low intensity conditions at Day-1 and Day-2. **B.** Group level beta rhythm suppression and rebound (n = 18) on both days and intensity conditions. Topographic maps show group level peak strengths of beta suppression and rebound in both intensity conditions. Note that beta suppression is the positive value calculated by the vector sum of the gradiometer pairs.

Fig. 5B illustrates group averaged (n = 18) spatial distribution of beta rhythm suppression and rebound on both days and intensity conditions. Beta rhythm suppression and rebound were observed after the stimulus onset in high and low intensity conditions. These responses were strongest in the contralateral hemisphere relative to the stimulated knee, particularly for beta rebound. As expected, the peak MEG-sensor pair for both suppression and rebound were localized over the sensorimotor cortex, with additional prominent beta suppression observed in more lateral sensor regions. Under high intensity condition, beta suppression peaked at approximately 280 ms following the stimulus onset, whereas under low intensity condition it peaked at ∼225 ms. Beta rebound was primarily localized to the contralateral SM1 cortex, peaking at ∼750 ms and ∼755 ms for both intensity conditions, respectively.

To further illustrate the cortical distribution of induced responses to proprioceptive stimulation, especially in beta suppression, source-level analysis was performed in a subset of four participants with available MRI data. Fig. 6A shows individual topographic maps in high and low intensity conditions at Day-1 and Day-2. Similar activation patterns were observed in the sensorimotor cortex following proprioceptive stimulation on both days and intensity conditions. As an illustrative example, the group-average map across these four participants during high intensity condition revealed that peak beta suppression activity was localized more towards the hand area of the SM1 cortex, while the peak beta rebound activity was in the knee representation area (Fig. 6B), aligning with the expected representation of the lower limb, although the sample size was limited.

**Fig. 6.**
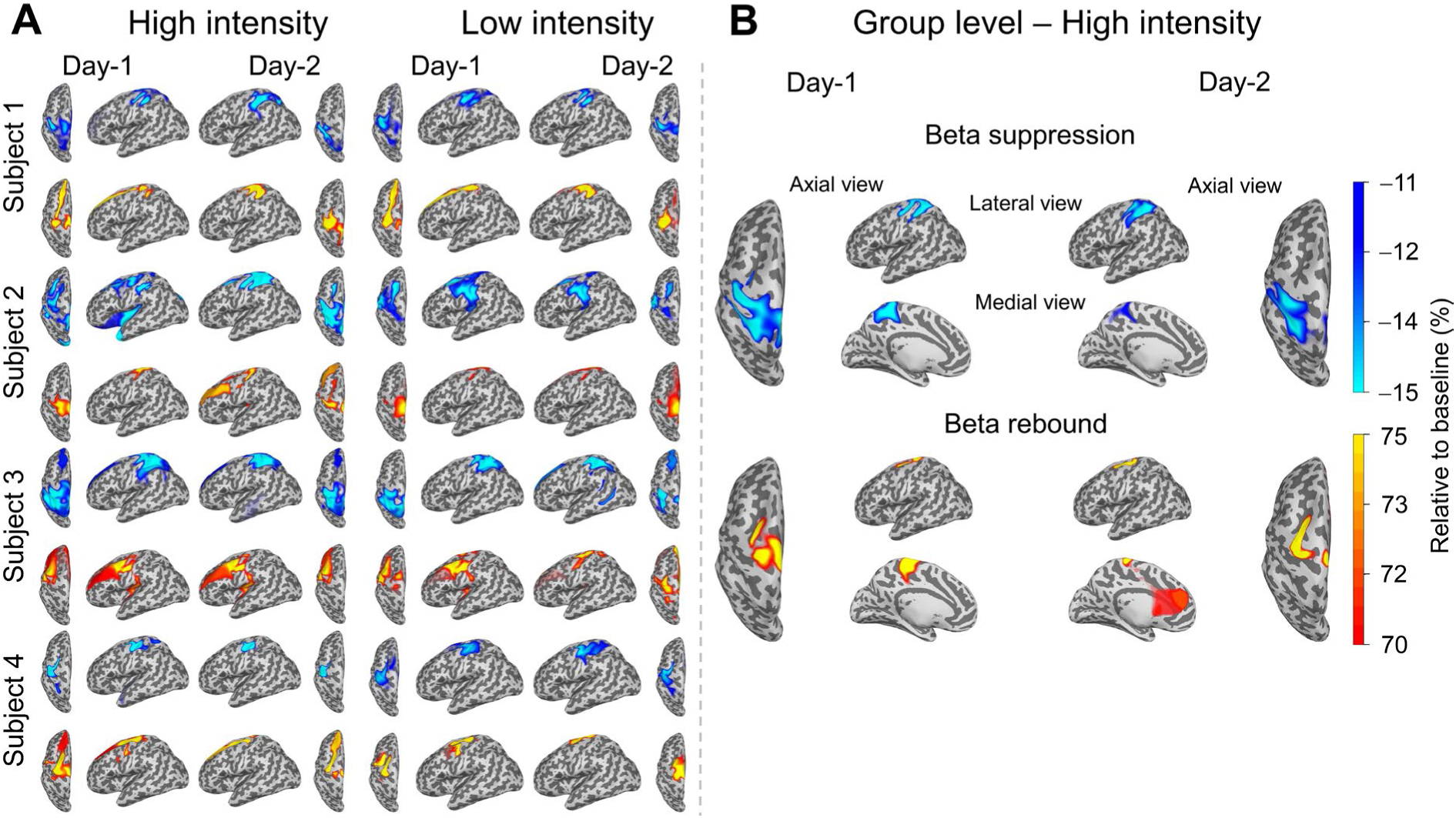
Induced responses at source-level to proprioceptive stimulation (n = 4). **A.** Individual source-level induced responses for 4 participants across both days and intensity conditions. The peak activity is shown in the contralateral (left) hemisphere to the stimulated knee. **B.** Group level induced responses projected on inflated brain for high intensity condition at Day-1 and Day-2.

### 3.6 Inter-session reliability

Fig. 7 illustrates the individual values of cortical and muscular responses, and acceleration magnitude and Fig. 8 shows their inter-session agreements between days. Overall, all variables were well repeatable under both intensity conditions for most of participants, especially for the cortical responses. As shown in the Bland-Altman plots (Fig. 8B), the cortical evoked and induced responses exhibited close agreement between days with narrow limits of bias. Intraclass correlation coefficient values indicated moderate to excellent reliability for both high (ICC: 0.89−0.96) and low intensity (ICC: 0.69−0.97) conditions. However, the ICC values appeared to be stronger in high than in low intensity conditions. Similarly, stronger correlations between Day-1 and Day-2 were observed in high intensity condition (r = 0.88−0.98).

**Fig. 7.**
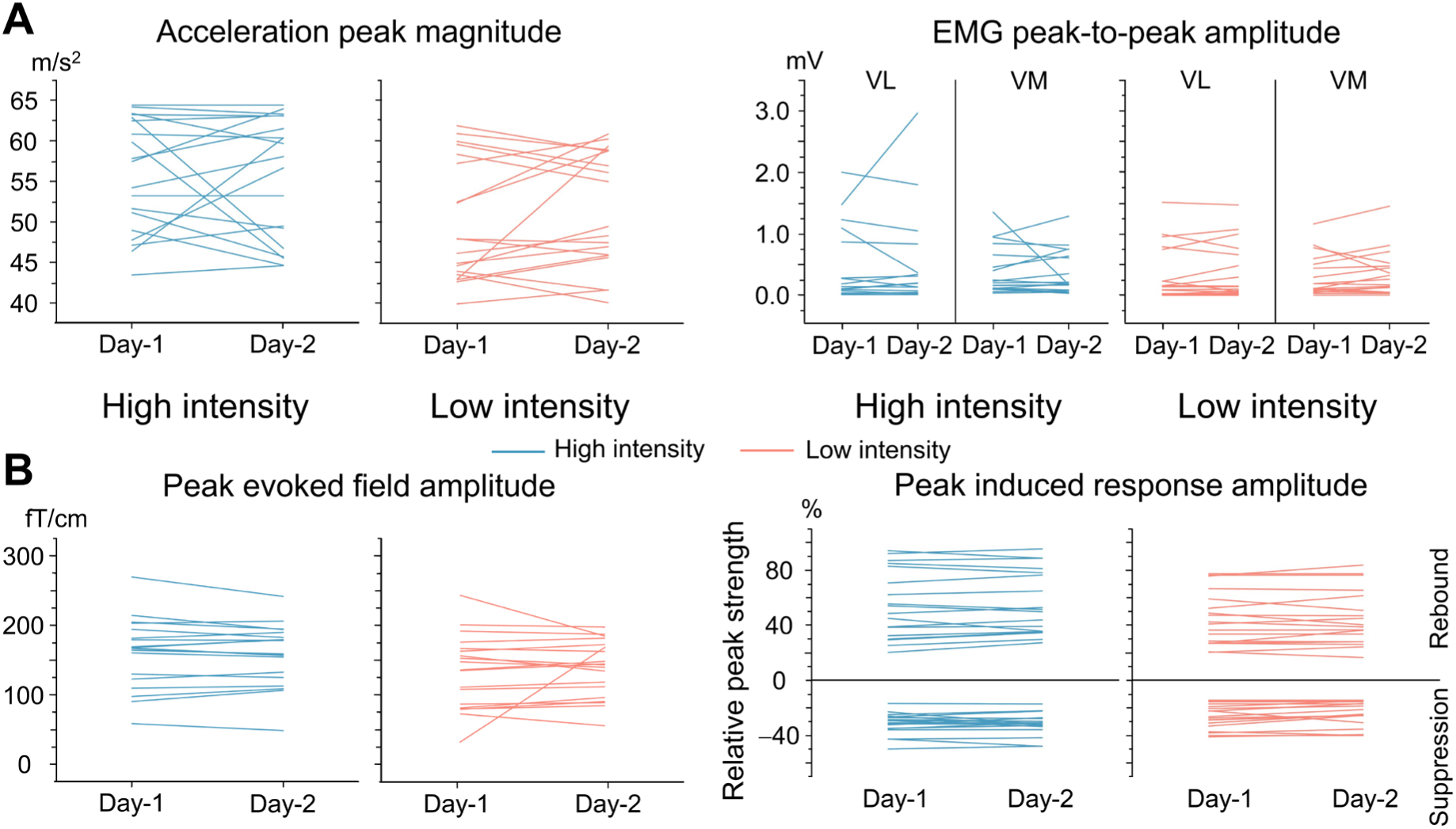
Individual subjects’ kinematics, muscular and cortical responses at Day-1 and Day-2 under both intensity conditions. Acceleration peak magnitude and EMG amplitudes (A), evoked and induced responses (B) at individual level between Day-1 and Day-2. Thin colorful lines represent direction of change for each participant separately in high (light blue) and low (light pink) intensity conditions.

**Fig. 8.**
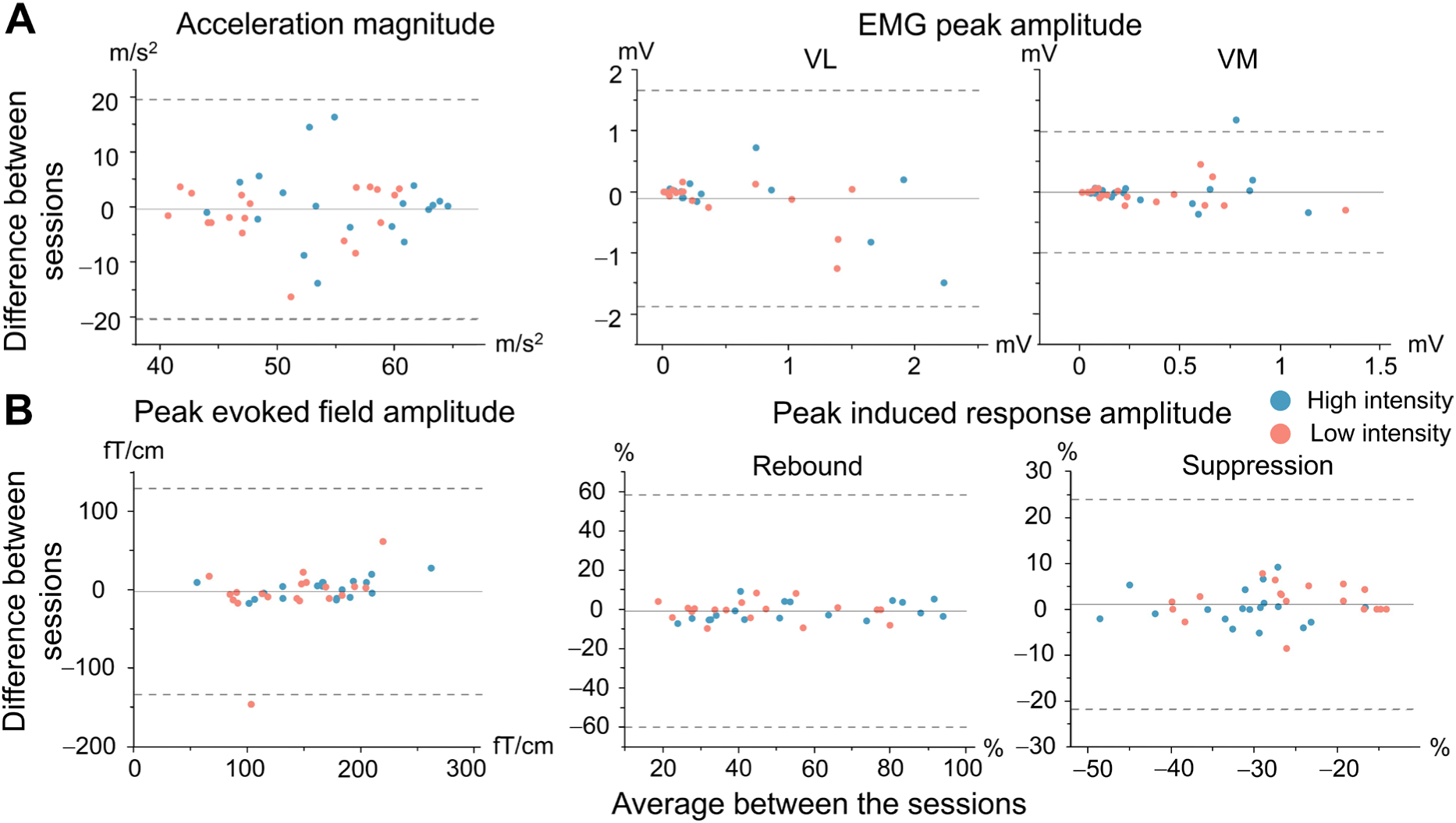
Intersession agreement for kinematics, muscular, evoked and induced cortical responses to high and low intensity conditions. **A.** Bland-Altman plots for kinematics and EMG (VL and VM muscles) between Day-1 and Day-2. **B.** Bland-Altman plots for evoked and induced (suppression and rebound) cortical responses between Day-1 and Day-2 separately. Solid black lines indicate the mean difference between days and dashed lines correspond to 95% confidence interval. The different colors indicate high (light blue) and low (light pink) intensity conditions, respectively.

Fig. 8A shows the Bland-Altman plots for muscular responses and acceleration magnitudes. Muscular responses in the VL and VM muscles were also reliable (ICC: 0.66−0.89), whereas acceleration magnitude showed small inter-session variation in high (Day-1: 55.8 ± 7.0 m/s^2^ vs. Day-2: 55.5 ± 7.5 m/s^2^) and low (Day-1: 50.0 ± 7.4 m/s^2^ vs. Day-2: 51.5 ± 7.2 m/s^2^) intensity conditions. More detailed reliability values are shown in Table 2.

**Table 2.** Intersession reliability of kinematics, muscular, evoked and induced responses for both high and low intensity conditions between Day-1 and Day-2.

|  | High intensity |  | Low intensity |  |
| --- | --- | --- | --- | --- |
|  | ICC | r | ICC | r |
| **EMG peak amplitude** |  |  |  |  |
| VL | 0.89 | 0.86*** | 0.80 | 0.87*** |
| VM | 0.66 | 0.65** | 0.88 | 0.88*** |
| **Evoked field** |  |  |  |  |
| Peak amplitude | 0.96 | 0.98*** | 0.69 | 0.72*** |
| **Induced response** |  |  |  |  |
| Peak suppression | 0.89 | 0.88*** | 0.90 | 0.91*** |
| Peak rebound | 0.96 | 0.98*** | 0.97 | 0.97*** |

Intraclass (ICC) and Pearson's (r) correlation coefficient values are presented in this table.
\*p < 0.05, \*\*p < 0.01, \*\*\*p < 0.001.
ICC < 0.5: poor; 0.5 < ICC < 0.75: moderate; 0.75 < ICC < 0.9: good; ICC > 0.9: excellent.

## 4. Discussion

We introduced a novel MEG-compatible device to stimulate specifically the proprioceptors of the knee extensors through patellar-tendon taps. Our results indicate that the stimulator can elicit robust cortical (MEG) and muscular (EMG) responses under two different intensity conditions (high and low). The tendon tapping was followed with typical stretch reflex in the quadriceps muscles and clear cortical responses in the SM1 cortex. The muscular and cortical responses induced by the novel stimulator showed moderate-to-excellent repeatability at the group level for both intensity conditions. For cortical responses, the repeatability level was in accordance with earlier evidence from finger or ankle joint proprioceptive stimulations using movement actuators in MEG (Illman et al., 2022; Mujunen et al., 2022). The cortical responses peaked in sensors above the contralateral leg region of the SM1 cortex, but surprisingly the beta suppression showed different spatial representation, peaking lateral to the conventional leg area, i.e., towards the hand region of the SM1 cortex, and was strong bilaterally. Overall, our observations prove that the novel stimulator is a feasible and reliable tool to be used in investigating the cortical processing of proprioceptive afference arising from the knee joint.

### 4.1 The repeatability of proprioceptive stimulation

The manual hammer-based stimulation was feasible and required only moderate training of the experimenter. The minor acoustic noise related to the hammer hitting the patellar tendon was well masked with the earplugs and background Brownian noise. The kinematics of the repeated stimuli were highly reliable from stimulus-to-stimulus, the CoV of peak acceleration magnitude was ∼2% within individual participants in both days and intensity conditions. Similarly, the session-to-session variation was on average 6% for high and 4% for low intensity condition at group level, as the peak acceleration magnitude of the tendon tap was practically the same between Day-1 and Day-2. Greater variability, ∼13% CoV, was observed from participant-to-participant variability, most likely due to small differences in the initial height and the accelerometer position, although the release height and accelerometer position were kept same between participants and sessions. To obtain more identical stimulation intensities between the participants, one should pay special attention to positioning the hammer and accelerometer at the same initial position, particularly ensuring consistent anterior-posterior alignment relative to the height controller across participants. Despite the above kinematic differences, the level of the stimulus variation was close to more controlled millisecond-accurate computer driven pneumatic movement actuators (i.e., proprioceptive stimulators) to evoke finger movements (Piitulainen et al., 2020; Piitulainen et al., 2018a) and ankle joint rotations (Chen et al., 2025; Mujunen et al., 2022; Piitulainen et al., 2018b). In conclusion, the knee-joint stimulator is a mechanically feasible and reliable tool to stimulate the proprioceptors through the patellar tendon in the MEG environment.

### 4.2 The muscular response to proprioceptive stimulation

Our results showed a typical stretch-reflex to proprioceptive stimulation in the knee extensors for both intensity conditions in majority of the participants. It is typical that the stretch reflex amplitude (EMG response) varies substantially from stimulus-to-stimulus. For example, the quadriceps stretch reflex amplitude has shown to vary ∼32% when evoked to resting muscle (Stam and Tan, 1987; Stam and van Crevel, 1989). In our case, the variation was ∼30%. Notably, the inter-session variation of the muscular response was also high on average about 30%, and in some participants up to 70–93%. The high inter-session variation could arise from differences in the participant’s state, such as mental or muscle fatigue (A et al., 2021; Moore et al., 2002). However, we scheduled the sessions to the same time of day for each participant to minimize circadian effects. In addition, the inter-individual variation of the muscular response was not calculated in this study, since the EMG differences can be largely due to anatomical differences of the muscle, subcutaneous tissue and muscle fiber innervation, which dramatically affects the amplitude of surface EMG (Farina et al., 2002). Due to these inherent sources of inter-individual variability, the absolute EMG responses cannot be meaningfully compared across individuals, and EMG normalization (e.g., MVC-based scaling) was not performed for tendon-tap reflex responses. Therefore, EMG results were interpreted only within subject and inter-session levels, rather than at the group inter-individual level.

### 4.3 The cortical responses to proprioceptive stimulation

To the best of our knowledge, this is the first study to investigate cortical evoked responses and beta modulations to tendon tap stimulation of the knee joint. We observed clear evoked and induced cortical responses in the SM1 cortex at both intensity conditions for all participants included in the final analysis. As expected, the kinematics of the stimulus did not correlate with the cortical evoked or induced response amplitudes in either intensity conditions. This means that variation in the cortical responses was not due to the small variation in the peak acceleration magnitude of the proprioceptive stimuli. Likewise, no statistically significant correlations were found between head height (position in z axis) and cortical response amplitudes in either evoked or induced responses (evoked field: p > 0.3; beta suppression: p > 0.67; beta rebound: p > 0.51), indicating that inevitable small (2 mm or less) difference in the head position between days did not systematically bias cortical responses.

The evoked fields remained stable between days with mean inter-session variation of ∼6%, which is much less compared to ∼30% inter-session variation in the muscular EMG response. Only three participants showed a higher inter-session variation with above 11%. The high repeatability of the evoked field likely stems from sufficient time-locked precision in the mechanical operation of the knee proprioceptive stimulator, ensuring stable afferent proprioceptive volley to the cortex. The repeatability was comparable to previously reported values for ankle-joint proprioceptive stimulation using mechanically robust pneumatic movement actuators (Mujunen et al., 2022). Thus, these findings demonstrate that the novel stimulator elicits highly repeatable evoked cortical responses, revealing that the novel stimulator can provide the reliable and time-locked proprioceptive input suitable for longitudinal MEG studies.

The beta rhythm suppression and rebound have been widely studied in somatosensory domains with different stimuli and tasks, including active movement (Espenhahn et al., 2017; Wilson et al., 2014), passive movement (Giangrande et al., 2024; Mujunen et al., 2025; Toledo et al., 2016; Walker et al., 2020) and tactile stimulation (Houdayer et al., 2006; Illman et al., 2022; Illman et al., 2020; Parkkonen et al., 2015). However, our study was the first one focusing on the naturalistic stimulation of the knee joint, providing novel observations in spatial domain of the beta rhythm modulation (see 4.4). In the amplitude domain, the knee-joint stimulation evoked a typical pattern of beta-band power modulation for both intensity conditions, characterized by prominent suppression followed by its rebound in good agreement with the previous studies on other joints or somatosensory stimuli.

The induced responses exhibited similar inter-session variability as the evoked fields, on average ∼7% for suppression and ∼6% for rebound. Few participants showed inter-session variability just above 20%, which is even less than what has been reported for proprioceptive stimulation of the hand or ankle joint using robust pneumatic movement actuators (Illman et al., 2022; Mujunen et al., 2022). However, the inter-individual variation was on average above 30% for beta suppression and 40% for beta rebound, which seems to be typical for the induced responses in MEG (Illman et al., 2022; Mujunen et al., 2022). The high inter-individual variability largely reflects differences in the functional anatomy of the cortex that affects the orientation of cortical currents relative to the MEG sensor orientation, and thus the MEG signal-to-noise ratio (Hämäläinen et al., 1993). It is noteworthy that the induced responses are much weaker than evoked fields, and thus are likely to be even more prone for the anatomical and other differences between the individuals. The representation for the lower limb in the primary somatosensory cortex lies within the paracentral lobules, deep in the interhemispheric fissure, being more challenging to detect with MEG compared, e.g., to the more superficial hand area (Jaeger et al., 2014; Kapreli et al., 2006; Piitulainen et al., 2015). In conclusion, the beta rhythm modulation is highly repeatable at the group level, but some caution is needed when following or interpreting individuals.

### 4.4 The spatial distribution of the cortical responses

For the evoked field, the MEG sensor showing the peak cortical response was approximately over the leg area of left SM1 cortex, as expected. Similar topography has been shown for ankle-joint stimuli (Chen et al., 2025; Mujunen et al., 2022; Piitulainen et al., 2018b), which have been shown to localize in the paracentral lobule corresponding to foot representation area (Mujunen et al., 2025). Notably, but not surprisingly, the topographies were systematically shifted slightly posteriorly due to the use of stabilizing foam cushion between the forehead and MEG-helmet during the MEG recordings.

Partly unexpected spatial activation pattern was observed for the induced responses, specifically for the beta suppression. The beta suppression peaked substantially more laterally towards the hand area when compared to the evoked field, i.e., beyond the typical leg region of the SM1 cortex, similar to what has been reported for proprioceptive beta suppression responses of the ankle joint (Mujunen et al., 2022). In addition, the suppression was strong bilaterally, although the suppression in other joints is typically clearly stronger in the hemisphere contralateral to the stimulus (Hari and Salmelin, 1997). Furthermore, the beta rebound peaked above the expected leg region. Because of this unexpected spatial pattern, we performed source-level analysis for those participants who had MRIs available (n = 4). Similar spatial patterns were confirmed at the source level. It thus seems that knee joint proprioception may have unique cortical representation compared to the more distal joints of the lower limb. Our observation may also be interpreted in light of recent work by Gordon et al. (2023), who identified inter-effector regions in the primary motor cortex that are interleaved between the classical hand, foot, and mouth representations. Importantly, movements requiring less precise motor control were shown to co-activate several inter-effector regions, whereas fine motor tasks of the hand or foot selectively activated their effector-specific areas. Indeed, knee-joint movements do not typically require precise fine motor control if compared to the more distal parts of the limbs, and thus it is plausible suggestion that the observed beta suppression localizes more to the inter-effector region.

In fMRI, wide cortical activations have been observed for the knee movement in the previous related studies. For example, Jaeger et al. (2014) investigated the brain activation between active and passive knee movements using fMRI. Their results demonstrated cortical activation in both SM1 cortex and the secondary somatosensory (SII) cortex during passive movements. A strong SII cortex activation has been reported for passive index-finger movements in fMRI when using pneumatic finger-movement actuator (Nurmi et al., 2018). The SII cortex is thought to participate in the initial processing of somatosensory input directly from thalamus and higher-order processing of input indirectly from the primary somatosensory cortex. In addition, the SII cortex receives somatosensory information from both sides of the body and typically shows bilateral response with unilateral stimulation (Ferretti et al., 2005; Li Hegner et al., 2007; Manzoni et al., 1986; Ridley and Ettlinger, 1976). Therefore, further studies are needed to confirm our novel observations beta rhythm modulation, especially for the beta suppression, to proprioceptive stimulation of the knee joint, and the role of SII cortex needs also to be clarified.

### 4.5 Advantages and limitations of the MEG-compatible knee-joint stimulator

Unlike actuator-based devices used in the previous proprioceptive MEG studies, such as pneumatic cylinder-lever system or DC-motor-based movement actuators, our stimulator delivered input through gravity-driven mechanical tapping of the patellar tendon. All aforementioned approaches are able to activate the proprioceptors by mechanically stimulating the muscles and tendon. The resulting afferent inputs split into two pathways: (i) spinal collaterals that synapse onto α-motoneuron pools and can evoke the monosynaptic stretch reflex, and (ii) the main ascending branch that continues through the dorsal column-medial lemniscus pathway, relays in thalamus, and reaches the cortex within ∼40 ms for the lower limb. It is important to note that the main cortical response is initiated by the first primary volley of the proprioceptive afference, whereas a secondary volley, arising from the reflexive muscle contraction when the stretch reflex is elicited, is delayed ∼60 ms with respect to the primary volley. The secondary volley can affect the later components of the cortical evoked and induced responses.

The mechanical configuration of our knee-joint stimulator ensured a stable gravity driven stimulation, and the stimulus intensity can be adjusted by varying the initial angle of the hammer, i.e., its “height”. A more critical challenge is to ensure the same position of the leg and the knee throughout the session and from day-to-day. To overcome this problem in future applications, the mechanical setup could be improved by incorporating a customized knee holder or adjustable brace to fix the knee joint angle and knee position in both anterior-posterior and medio-lateral directions more precisely. Nevertheless, the tapping intensity remained practically constant during intensity conditions and days. According to Marshall and Little (2002), the most significant factor which could influence the tendon reflex response is the tapping intensity to the tendon. Stable stimulation is crucial for future longitudinal MEG studies of proprioceptive afference, but fortunately small variation in the stimulus intensity did not affect significantly the cortical responses, which is in line with studies showing that movement stimulation parameters, such as range of the evoked movement, does not significantly affect the cortical response amplitude (Nurmi et al., 2023).

Proprioceptive stimulation inducing stretch-reflex may cause delayed mechanical body and head movements through the sudden strong activation of the stimulated muscles, especially in participants with low stretch reflex threshold. In one of our participants, with exceptionally sensitive and strong stretch reflex (i.e. strong activation of the knee extensors), some head movement-related artefacts were observed occasionally in the MEG signals. Thus, we had to exclude this participant from the final analysis. It is possible that such a strong stretch reflex may also introduce mechanical vibration of MEG chair, which contains some metallic parts that may cause artifacts directly to the MEG signals, but are expected to be well removed by Maxfilter since the magnetic source is far from the MEG helmet head space. Nevertheless, we recommend using foam cushions between the head and the helmet to minimize possible head movements. Use of adjustable straps to secure the leg and body firmly to the chair during the stimulation is also recommended to maintain the knee and the stimulation site at fixed position.

The stimulating hammer was manually released by the experimenter. A fully motorized or electronically triggered release would be more precise but would also omit the need for the presence of extra person in the shielded room. Future developments should therefore aim to implement an automated release mechanism to further minimize manual user dependence. Nevertheless, these limitations are unlikely to compromise the main conclusions of the present study regarding the feasibility and reliability of the knee-joint stimulator.

### 4.6 Study limitations and future perspectives

Some methodological limitations should be considered in future experiments attempting to use same or similar stimulation approach. First, our stimulation intensities were the same for each participant, but the stretch-reflex threshold varies between the individuals. Optimally, one would first define the stretch-reflex threshold, and then would use intensity relative to it for more individualized approach. In addition, attentional state was not systematically controlled, so their potential contribution to the recorded cortical signals cannot be fully excluded (Piitulainen et al., 2021). The generalizability of the current data is limited, since we recruited only healthy young adults, which limits the application to patient populations, where reflex characteristics and cortical responsiveness may differ. Finally, only four of our participants had anatomical MRI available, which restricted our conclusions from the source level analysis when interpreting the beta suppression localization. The source-level analysis was not planned a priori, but was done after we observed the unexpected spatial distribution of beta suppression, and thus structural MRI for the full sample was not possible. Thus, our main conclusions regarding feasibility and reliability are based primarily on sensor-level analyses, and the anatomical interpretations should be treated with caution.

Additionally, although we provided the results of high and low intensity conditions, the comparison between different intensity conditions was not completed in this study. The novel stimulator could also be used below stretch-reflex intensity, as in previous actuator-based studies of passive joint rotations (Nurmi et al., 2018; Piitulainen et al., 2015; Zhao et al., 2025), to examine how stimulation strength shapes the magnitude and spatial patterns of cortical proprioceptive responses.

## 5. Conclusion

Our study demonstrated that the novel MEG-compatible stimulator is a feasible and reliable tool to be used in quantifying and following the cortical processing of proprioceptive afference from the knee extensors. The cortical evoked and induced and muscular stretch-reflex responses elicited by the novel stimulator were consistent from day-to-day. The cortical responses peaked at MEG sensors over the expected contralateral leg area of SM1 cortex, with the exception of beta suppression peaking lateral to the expected leg region, i.e., towards the hand area, being prominent bilaterally. This novel observation seems to suggest unique representation for beta suppression in the knee joint. The novel stimulator has the potential to investigate and follow knee joint proprioception in various patients and interventions at the group level, such as in elderly and patients with knee osteoarthritis, total-knee replacement or anterior cruciate ligament injury, but caution is needed when assessing individual participants.

## Reference

A, D.G., Teodosio, C., Pezarat-Correia, P., Vila-Cha, C., G, V.M., 2021. Effects of acute sleep deprivation on H reflex and V wave. J Sleep Res 30, e13118.10.1111/jsr.13118.

Alary, F., Simoes, C., Jousmaki, V., Forss, N., Hari, R., 2002. Cortical activation associated with passive movements of the human index finger: an MEG study. Neuroimage 15, 691–696.10.1006/nimg.2001.1010.

Banios, K., Raoulis, V., Fyllos, A., Chytas, D., Mitrousias, V., Zibis, A., 2022. Anterior and Posterior Cruciate Ligaments Mechanoreceptors: A Review of Basic Science. Diagnostics (Basel) 12.10.3390/diagnostics12020331.

Barbero, M., Merletti, R., Rainoldi, A., 2012. Atlas of muscle innervation zones : understanding surface electromyography and its applications. Springer, Milan; New York.

Barrett, D.S., Cobb, A.G., Bentley, G., 1991. Joint proprioception in normal, osteoarthritic and replaced knees. J Bone Joint Surg Br 73, 53–56.10.1302/0301-620X.73B1.1991775.

Bourguignon, M., De Tiege, X., Op de Beeck, M., Pirotte, B., Van Bogaert, P., Goldman, S., Hari, R., Jousmaki, V., 2011. Functional motor-cortex mapping using corticokinematic coherence. Neuroimage 55, 1475–1479.10.1016/j.neuroimage.2011.01.031.

Bourguignon, M., Piitulainen, H., De Tiege, X., Jousmaki, V., Hari, R., 2015. Corticokinematic coherence mainly reflects movement-induced proprioceptive feedback. Neuroimage 106, 382–390.10.1016/j.neuroimage.2014.11.026.

Chen, J., Mujunen, T., Piitulainen, H., 2025. Reproducibility of corticokinematic coherence for proprioceptive stimulation of the ankle joint. Neuroscience 589, 230–239.10.1016/j.neuroscience.2025.10.049.

Cheyne, D., Bakhtazad, L., Gaetz, W., 2006. Spatiotemporal mapping of cortical activity accompanying voluntary movements using an event-related beamforming approach. Hum Brain Mapp 27, 213–229.10.1002/hbm.20178.

Daneshjoo, A., Mokhtar, A.H., Rahnama, N., Yusof, A., 2012. The effects of comprehensive warm-up programs on proprioception, static and dynamic balance on male soccer players. PLoS One 7, e51568.10.1371/journal.pone.0051568.

David, O., Kilner, J.M., Friston, K.J., 2006. Mechanisms of evoked and induced responses in MEG/EEG. Neuroimage 31, 1580–1591.10.1016/j.neuroimage.2006.02.034.

Druschky, K., Kaltenhauser, M., Hummel, C., Druschky, A., Huk, W.J., Neundorfer, B., Stefan, H., 2003. Somatosensory evoked magnetic fields following passive movement compared with tactile stimulation of the index finger. Exp Brain Res 148, 186–195.10.1007/s00221-002-1293-4.

Elias, L.J., Bryden, M.P., Bulman-Fleming, M.B., 1998. Footedness is a better predictor than is handedness of emotional lateralization. Neuropsychologia 36, 37–43.10.1016/s0028-3932(97)00107-3.

Espenhahn, S., de Berker, A.O., van Wijk, B.C.M., Rossiter, H.E., Ward, N.S., 2017. Movement-related beta oscillations show high intra-individual reliability. Neuroimage 147, 175–185.10.1016/j.neuroimage.2016.12.025.

Farina, D., Cescon, C., Merletti, R., 2002. Influence of anatomical, physical, and detection-system parameters on surface EMG. Biol Cybern 86, 445–456.10.1007/s00422-002-0309-2.

Ferretti, A., Arienzo, D., Del Gratta, C., Caulo, M., Babiloni, C., Tartaro, A., Rossini, P., Romani, G., 2005. A BOLD-fMRI study of the response in primary and secondary somatosensory cortices elicited by electric median nerve stimulation at different frequencies. IJBEM 7, 1–3

Gaetz, W., Cheyne, D., 2006. Localization of sensorimotor cortical rhythms induced by tactile stimulation using spatially filtered MEG. Neuroimage 30, 899–908.10.1016/j.neuroimage.2005.10.009.

Giangrande, A., Mujunen, T., Luigi Cerone, G., Botter, A., Piitulainen, H., 2024. Maintained volitional activation of the muscle alters the cortical processing of proprioceptive afference from the ankle joint. Neuroscience 560, 314–325.10.1016/j.neuroscience.2024.09.049.

Gordon, E.M., Chauvin, R.J., Van, A.N., Rajesh, A., Nielsen, A., Newbold, D.J., Lynch, C.J., Seider, N.A., Krimmel, S.R., Scheidter, K.M., Monk, J., Miller, R.L., Metoki, A., Montez, D.F., Zheng, A., Elbau, I., Madison, T., Nishino, T., Myers, M.J., Kaplan, S., Badke D’Andrea, C., Demeter, D.V., Feigelis, M., Ramirez, J.S.B., Xu, T., Barch, D.M., Smyser, C.D., Rogers, C.E., Zimmermann, J., Botteron, K.N., Pruett, J.R., Willie, J.T., Brunner, P., Shimony, J.S., Kay, B.P., Marek, S., Norris, S.A., Gratton, C., Sylvester, C.M., Power, J.D., Liston, C., Greene, D.J., Roland, J.L., Petersen, S.E., Raichle, M.E., Laumann, T.O., Fair, D.A., Dosenbach, N.U.F., 2023. A somato-cognitive action network alternates with effector regions in motor cortex. Nature 617, 351–359.10.1038/s41586-023-05964-2.

Gramfort, A., Luessi, M., Larson, E., Engemann, D.A., Strohmeier, D., Brodbeck, C., Parkkonen, L., Hamalainen, M.S., 2014. MNE software for processing MEG and EEG data. Neuroimage 86, 446–460.10.1016/j.neuroimage.2013.10.027.

Hämäläinen, M., Hari, R., Ilmoniemi, R.J., Knuutila, J., Lounasmaa, O.V., 1993. Magnetoencephalography---theory, instrumentation, and applications to noninvasive studies of the working human brain. Reviews of Modern Physics 65, 413–497.10.1103/RevModPhys.65.413.

Hari, R., Salmelin, R., 1997. Human cortical oscillations: a neuromagnetic view through the skull. Trends in Neurosciences 20, 44–49.10.1016/S0166-2236(96)10065-5.

Heinrichs-Graham, E., Kurz, M.J., Becker, K.M., Santamaria, P.M., Gendelman, H.E., Wilson, T.W., 2014. Hypersynchrony despite pathologically reduced beta oscillations in patients with Parkinson’s disease: a pharmaco-magnetoencephalography study. J Neurophysiol 112, 1739–1747.10.1152/jn.00383.2014.

Hollnagel, C., Brugger, M., Vallery, H., Wolf, P., Dietz, V., Kollias, S., Riener, R., 2011. Brain activity during stepping: a novel MRI-compatible device. J Neurosci Methods 201, 124–130.10.1016/j.jneumeth.2011.07.022.

Houdayer, E., Labyt, E., Cassim, F., Bourriez, J.L., Derambure, P., 2006. Relationship between event-related beta synchronization and afferent inputs: analysis of finger movement and peripheral nerve stimulations. Clin Neurophysiol 117, 628–636.10.1016/j.clinph.2005.12.001.

Hyvarinen, A., Oja, E., 2000. Independent component analysis: algorithms and applications. Neural Netw 13, 411–430.10.1016/s0893-6080(00)00026-5.

Illman, M., Laaksonen, K., Jousmaki, V., Forss, N., Piitulainen, H., 2022. Reproducibility of Rolandic beta rhythm modulation in MEG and EEG. J Neurophysiol 127, 559–570.10.1152/jn.00267.2021.

Illman, M., Laaksonen, K., Liljestrom, M., Jousmaki, V., Piitulainen, H., Forss, N., 2020. Comparing MEG and EEG in detecting the ∼20-Hz rhythm modulation to tactile and proprioceptive stimulation. Neuroimage 215, 116804.10.1016/j.neuroimage.2020.116804.

Jaeger, L., Marchal-Crespo, L., Wolf, P., Riener, R., Michels, L., Kollias, S., 2014. Brain activation associated with active and passive lower limb stepping. Front Hum Neurosci 8, 828.10.3389/fnhum.2014.00828.

Jurkiewicz, M.T., Gaetz, W.C., Bostan, A.C., Cheyne, D., 2006. Post-movement beta rebound is generated in motor cortex: evidence from neuromagnetic recordings. Neuroimage 32, 1281–1289.10.1016/j.neuroimage.2006.06.005.

Kapreli, E., Athanasopoulos, S., Papathanasiou, M., Van Hecke, P., Strimpakos, N., Gouliamos, A., Peeters, R., Sunaert, S., 2006. Lateralization of brain activity during lower limb joints movement. An fMRI study. Neuroimage 32, 1709–1721.10.1016/j.neuroimage.2006.05.043.

Koo, T.K., Li, M.Y., 2016. A Guideline of Selecting and Reporting Intraclass Correlation Coefficients for Reliability Research. J Chiropr Med 15, 155–163.10.1016/j.jcm.2016.02.012.

Koralewicz, L.M., Engh, G.A., 2000. Comparison of proprioception in arthritic and age-matched normal knees. J Bone Joint Surg Am 82, 1582–1588.10.2106/00004623-200011000-00011.

Laaksonen, K., Kirveskari, E., Makela, J.P., Kaste, M., Mustanoja, S., Nummenmaa, L., Tatlisumak, T., Forss, N., 2012. Effect of afferent input on motor cortex excitability during stroke recovery. Clin Neurophysiol 123, 2429–2436.10.1016/j.clinph.2012.05.017.

Lakatos, P., O’Connell, M.N., Barczak, A., Mills, A., Javitt, D.C., Schroeder, C.E., 2009. The leading sense: supramodal control of neurophysiological context by attention. Neuron 64, 419–430.10.1016/j.neuron.2009.10.014.

Lange, R., Nowak, H., Haueisen, J., Weiller, C., 2001. Passive finger movement evoked fields in magnetoencephalography. Exp Brain Res 136, 194–199.10.1007/s002210000581.

Li Hegner, Y., Saur, R., Veit, R., Butts, R., Leiberg, S., Grodd, W., Braun, C., 2007. BOLD adaptation in vibrotactile stimulation: neuronal networks involved in frequency discrimination. J Neurophysiol 97, 264–271.10.1152/jn.00617.2006.

Manzoni, T., Conti, F., Fabri, M., 1986. Callosal projections from area SII to SI in monkeys: anatomical organization and comparison with association projections. J Comp Neurol 252, 245–263.10.1002/cne.902520208.

Marshall, G.L., Little, J.W., 2002. Deep tendon reflexes: a study of quantitative methods. J Spinal Cord Med 25, 94–99.10.1080/10790268.2002.11753608.

Martinez, M., Villagra, F., Loayza, F., Vidorreta, M., Arrondo, G., Luis, E., Diaz, J., Echeverria, M., Fernandez-Seara, M.A., Pastor, M.A., 2014. MRI-compatible device for examining brain activation related to stepping. IEEE Trans Med Imaging 33, 1044–1053.10.1109/TMI.2014.2301493.

Moore, B.D., Drouin, J., Gansneder, B.M., Shultz, S.J., 2002. The differential effects of fatigue on reflex response timing and amplitude in males and females. J Electromyogr Kinesiol 12, 351–360.10.1016/s1050-6411(02)00032-9.

Mujunen, T., Seipäjärvi, S., Nurminen, M., Parviainen, T., Piitulainen, H., 2022. Reproducibility of evoked and induced MEG responses to proprioceptive stimulation of the ankle joint. Neuroimage: Reports 2, 100110.10.1016/j.ynirp.2022.100110.

Mujunen, T., Sompa, U., Munoz-Ruiz, M., Monto, E., Rissanen, V., Ruuskanen, H., Ahtiainen, P., Piitulainen, H., 2025. Early peripheral nerve impairments in type 1 diabetes are associated with cortical inhibition of ankle joint proprioceptive afference. Clin Neurophysiol 173, 99–112.10.1016/j.clinph.2025.02.277.

Neuper, C., Wortz, M., Pfurtscheller, G., 2006. ERD/ERS patterns reflecting sensorimotor activation and deactivation. Prog Brain Res 159, 211–222.10.1016/S0079-6123(06)59014-4.

Nurmi, T., Hakonen, M., Bourguignon, M., Piitulainen, H., 2023. Proprioceptive response strength in the primary sensorimotor cortex is invariant to the range of finger movement. Neuroimage 269, 119937.10.1016/j.neuroimage.2023.119937.

Nurmi, T., Henriksson, L., Piitulainen, H., 2018. Optimization of Proprioceptive Stimulation Frequency and Movement Range for fMRI. Front Hum Neurosci 12, 477.10.3389/fnhum.2018.00477.

Nyland, J., Gamble, C., Franklin, T., Caborn, D.N.M., 2017. Permanent knee sensorimotor system changes following ACL injury and surgery. Knee Surg Sports Traumatol Arthrosc 25, 1461–1474.10.1007/s00167-017-4432-y.

Parkkonen, E., Laaksonen, K., Piitulainen, H., Parkkonen, L., Forss, N., 2015. Modulation of the reverse similar20-Hz motor-cortex rhythm to passive movement and tactile stimulation. Brain Behav 5, e00328.10.1002/brb3.328.

Pascual-Marqui, R.D., Lehmann, D., Koukkou, M., Kochi, K., Anderer, P., Saletu, B., Tanaka, H., Hirata, K., John, E.R., Prichep, L., Biscay-Lirio, R., Kinoshita, T., 2011. Assessing interactions in the brain with exact low-resolution electromagnetic tomography. Philos Trans A Math Phys Eng Sci 369, 3768–3784.10.1098/rsta.2011.0081.

Pfurtscheller, G., 2001. Functional brain imaging based on ERD/ERS. Vision Res 41, 1257–1260.10.1016/s0042-6989(00)00235-2.

Piitulainen, H., Bourguignon, M., De Tiege, X., Hari, R., Jousmaki, V., 2013. Corticokinematic coherence during active and passive finger movements. Neuroscience 238, 361–370.10.1016/j.neuroscience.2013.02.002.

Piitulainen, H., Bourguignon, M., Hari, R., Jousmaki, V., 2015. MEG-compatible pneumatic stimulator to elicit passive finger and toe movements. Neuroimage 112, 310–317.10.1016/j.neuroimage.2015.03.006.

Piitulainen, H., Illman, M., Jousmaki, V., Bourguignon, M., 2020. Feasibility and reproducibility of electroencephalography-based corticokinematic coherence. J Neurophysiol 124, 1959–1967.10.1152/jn.00562.2020.

Piitulainen, H., Illman, M., Laaksonen, K., Jousmaki, V., Forss, N., 2018a. Reproducibility of corticokinematic coherence. Neuroimage 179, 596–603.10.1016/j.neuroimage.2018.06.078.

Piitulainen, H., Nurmi, T., Hakonen, M., 2021. Attention directed to proprioceptive stimulation alters its cortical processing in the primary sensorimotor cortex. Eur J Neurosci.10.1111/ejn.15251.

Piitulainen, H., Seipajarvi, S., Avela, J., Parviainen, T., Walker, S., 2018b. Cortical Proprioceptive Processing Is Altered by Aging. Front Aging Neurosci 10, 147.10.3389/fnagi.2018.00147.

Ridley, R.M., Ettlinger, G., 1976. Impaired tactile learning and retention after removals of the second somatic sensory projection cortex (SII) in the monkey. Brain Res 109, 656–660.10.1016/0006-8993(76)90048-2.

Rossiter, H.E., Boudrias, M.H., Ward, N.S., 2014a. Do movement-related beta oscillations change after stroke? J Neurophysiol 112, 2053–2058.10.1152/jn.00345.2014.

Rossiter, H.E., Davis, E.M., Clark, E.V., Boudrias, M.H., Ward, N.S., 2014b. Beta oscillations reflect changes in motor cortex inhibition in healthy ageing. Neuroimage 91, 360–365.10.1016/j.neuroimage.2014.01.012.

Salmelin, R., Hari, R., 1994. Spatiotemporal characteristics of sensorimotor neuromagnetic rhythms related to thumb movement. Neuroscience 60, 537–550.10.1016/0306-4522(94)90263-1.

Solomonow, M., Krogsgaard, M., 2001. Sensorimotor control of knee stability. A review. Scand J Med Sci Sports 11, 64–80.10.1034/j.1600-0838.2001.011002064.x.

Stam, J., Tan, K.M., 1987. Tendon reflex variability and method of stimulation. Electroencephalogr Clin Neurophysiol 67, 463–467.10.1016/0013-4694(87)90010-1.

Stam, J., van Crevel, H., 1989. Measurement of tendon reflexes by surface electromyography in normal subjects. J Neurol 236, 231–237.10.1007/BF00314505.

Taulu, S., Simola, J., 2006. Spatiotemporal signal space separation method for rejecting nearby interference in MEG measurements. Phys Med Biol 51, 1759–1768.10.1088/0031-9155/51/7/008.

Toledo, D.R., Barela, J.A., Manzano, G.M., Kohn, A.F., 2016. Age-related differences in EEG beta activity during an assessment of ankle proprioception. Neurosci Lett 622, 1–5.10.1016/j.neulet.2016.04.028.

Vinding, M.C., Tsitsi, P., Piitulainen, H., Waldthaler, J., Jousmaki, V., Ingvar, M., Svenningsson, P., Lundqvist, D., 2019. Attenuated beta rebound to proprioceptive afferent feedback in Parkinson’s disease. Sci Rep 9, 2604.10.1038/s41598-019-39204-3.

Walker, S., Monto, S., Piirainen, J.M., Avela, J., Tarkka, I.M., Parviainen, T.M., Piitulainen, H., 2020. Older Age Increases the Amplitude of Muscle Stretch-Induced Cortical Beta-Band Suppression But Does not Affect Rebound Strength. Front Aging Neurosci 12, 117.10.3389/fnagi.2020.00117.

Wilson, T.W., Heinrichs-Graham, E., Becker, K.M., 2014. Circadian modulation of motor-related beta oscillatory responses. Neuroimage 102 Pt 2, 531–539.10.1016/j.neuroimage.2014.08.013.

Zhao, S., Bao, Z., Shou, X., 2025. Investigation of Proprioceptive Responses to Passive Lower Limb Movements Based on Corticokinematic Coherence. IEEE Trans Neural Syst Rehabil Eng 33, 2556–2564.10.1109/TNSRE.2025.3583668.

